# Mutational signatures associated with tobacco smoking in human cancer

**DOI:** 10.1101/051417

**Authors:** Ludmil B. Alexandrov, Young Seok Ju, Kerstin Haase, Peter Van Loo, Iñigo Martincorena, Serena Nik-Zainal, Yasushi Totoki, Akihiro Fujimoto, Hidewaki Nakagawa, Tatsuhiro Shibata, Peter J. Campbell, Paolo Vineis, David H. Phillips, Michael R. Stratton

**Affiliations:** Theoretical Biology and Biophysics (T-6), Los Alamos National Laboratory, Los Alamos, NM 87545, USA; Center for Nonlinear Studies, Los Alamos National Laboratory, Los Alamos, NM 87545, USA; University of New Mexico Comprehensive Cancer Center, Albuquerque, NM 87102, USA; Graduate School of Medical Science and Engineering (GSMSE), Korea Advanced Institute of Science and Technology (KAIST), Daejeon 34141, Republic of Korea; The Francis Crick Institute, Lincoln's Inn Fields Laboratory, 44 Lincoln's Inn Fields, London WC2A 3LY; Human Genome Laboratory, Department of Human Genetics, University of Leuven, 3000 Leuven, Belgium; Cancer Genome Project, Wellcome Trust Sanger Institute, Hinxton CB10 1SA, Cambridgeshire, UK; Department of Medical Genetics, Addenbrooke’s Hospital National Health Service (NHS) Trust, Cambridge, UK; Division of Cancer Genomics, National Cancer Center Research Institute, Chuo-ku, Tokyo, Japan.; Laboratory for Genome Sequencing Analysis, RIKEN Center for Integrative Medical Sciences, Tokyo, Japan; Department of Drug Discovery Medicine, Kyoto University Graduate School of Medicine, Kyoto, 606-8507, Japan; Laboratory of Molecular Medicine, Human Genome Center, The Institute of Medical Science, The University of Tokyo, Minato-ku, Tokyo, Japan; Department of Haematology, University of Cambridge, Cambridge CB2 0XY, UK; Human Genetics Foundation (HuGeF), 10126 Torino, Italy; Department of Epidemiology and Biostatistics, MRC-PHE Centre for Environment and Health, School of Public Health, Imperial College London, Norfolk Place, London W2 1PG, UK; King’s College London, MRC-PHE Centre for Environment and Health, Analytical and Environmental Sciences Division, Franklin-Wilkins Building, 150 Stamford Street, London SE1 9NH, UK

## Abstract

Tobacco smoking increases the risk of at least 15 classes of cancer. We analyzed somatic mutations and DNA methylation in 5,243 cancers of types for which tobacco smoking confers an elevated risk. Smoking is associated with increased mutation burdens of multiple distinct mutational signatures, which contribute to different extents in different cancers. One of these signatures, mainly found in cancers derived from tissues directly exposed to tobacco smoke, is attributable to misreplication of DNA damage caused by tobacco carcinogens. Others likely reflect indirect activation of DNA editing by APOBEC cytidine deaminases and of an endogenous clock-like mutational process. The results are consistent with the proposition that smoking increases cancer risk by increasing the somatic mutation load, although direct evidence for this mechanism is lacking in some smoking-related cancer types.

**ONE SENTENCE SUMMARY:** Multiple distinct mutational processes associate with tobacco smoking in cancer reflecting direct and indirect effects of tobacco smoke.

## MAIN TEXT

Exposure to tobacco smoke is a major cause of cancer worldwide, claiming the lives of more than six million people every year *(1–3)*. Tobacco smoking has been epidemiologically associated with cancers of the lung, larynx, naso/oro/hypopharynx, oral/nasal cavities, esophagus, bladder, liver, stomach, bone marrow (myeloid leukemia), ovary, uterine cervix, kidney, pancreas and colorectum *(4)* (Table 1). Tobacco smoke is a complex mixture of organic and inorganic chemicals among which at least 60 are carcinogens *(5)*. Many of these carcinogens are thought to cause cancer by inducing DNA damage which, if misreplicated, leads to an increased burden of somatic mutations and hence an elevated chance of acquiring “driver” mutations in cancer genes. Such damage is often in the form of covalent bonding of metabolically activated reactive species of the carcinogen to DNA bases, termed “DNA adducts” *(6)*. Tissues directly exposed to tobacco smoke (lung, larynx, pharynx, oral and nasal tissue) as well as some tissues not directly exposed (*e.g.*, bladder and cervix) show elevated levels of DNA adducts in smokers and thus evidence of exposure to carcinogenic components of tobacco smoke *(7, 8)*.

**Table 1.**
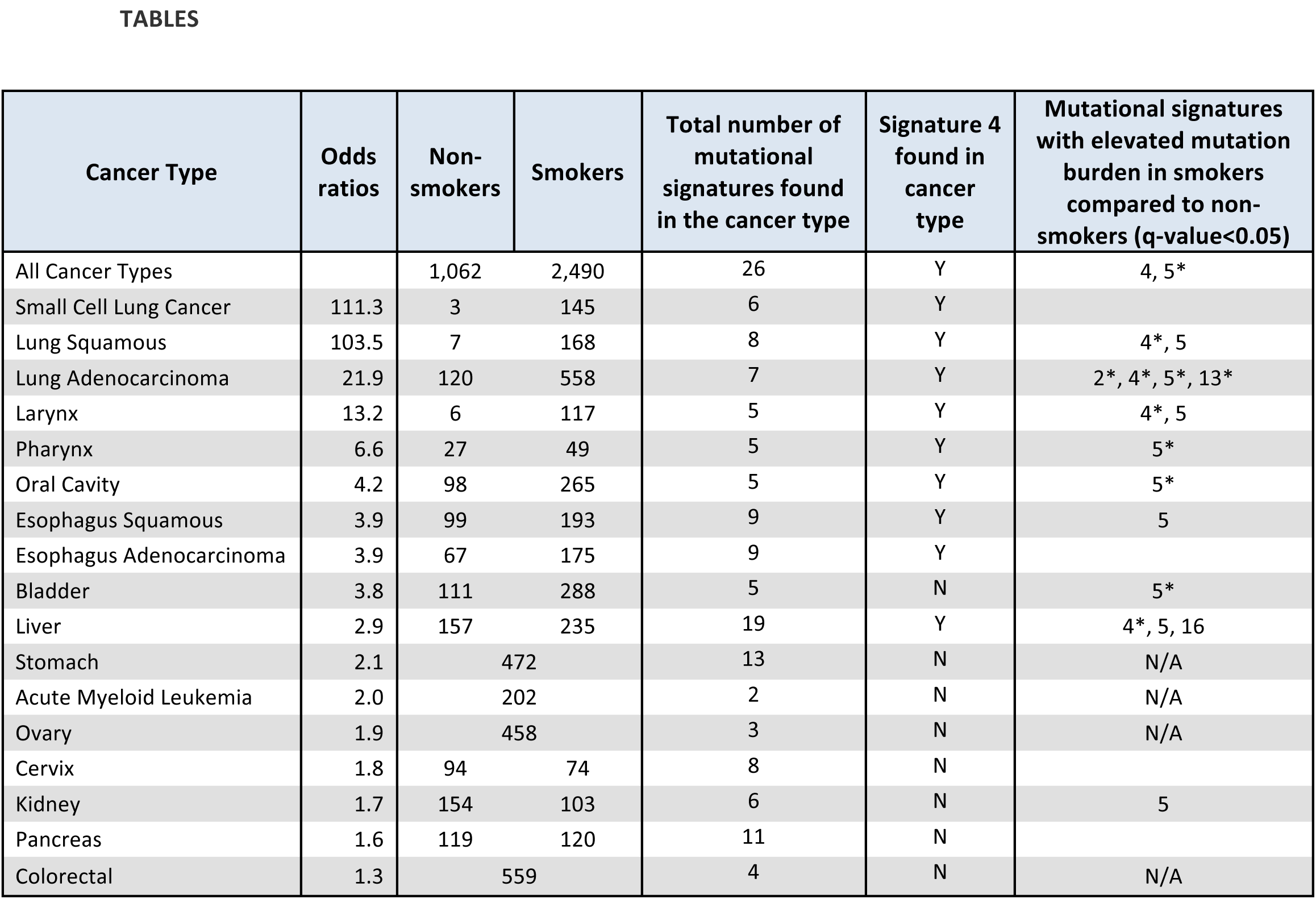
Mutational signatures and cancer types associated with tobacco smoking. Information about the age adjusted odds ratios for current male smokers to develop cancer is taken from refs. *(2, 3)*. Odds ratios for small cell lung cancer, lung squamous and lung adenocarcinoma are for an average daily dose of more than 30 cigarettes. Odds ratios for cervix and ovary are for current female smokers. Detailed information about all mutation types, all mutational signatures and DNA methylation is provided in Table S2. Nomenclature for signature IDs is consistent with the COSMIC website, http://cancer.sanger.ac.uk/cosmic/signatures. The patterns of all mutational signatures with elevated mutation burden in smokers are displayed in Fig. 2. N/A denotes lack of smoking annotation for a give cancer type. * denotes that a signature correlates with pack years smoked in a cancer type.

Each biological process causing mutations in the genomes of somatic cells leaves a characteristic mutational signature *(9)*. Many human cancers have a somatic mutation in the *TP53* gene and thus catalogues of *TP53* mutations compiled two decades ago enabled early exploration of these signatures *(10)*. The *TP53* mutation spectrum in lung cancers from smokers is different from that of non-smokers with a higher frequency of C>A transversions, commonly at methylated CpG sites *(11–14)*. To investigate mutational signatures using the thousands of catalogues of somatic mutations generated by systematic cancer exome and whole genome sequencing, we recently described an analytic framework in which each base substitution mutational signature is characterized using a 96 subclass mutation classification that includes the six substitution types C>A, C>G, C>T, T>A, T>C and T>G (all base substitutions are referred to by the pyrimidine of the Watson-Crick base pair) together with the bases immediately 5′ and 3′ to the mutated base *(15)*. The analysis extracts mutational signatures from series of somatic mutation catalogues and estimates the number of mutations contributed by each signature to each cancer genome *(15)*. Using this approach, more than 30 different base substitution mutational signatures have been identified across human cancer *(16–18)*.

Here, we studied 5,243 cancer genome sequences (4,633 exomes and 610 whole genomes curated from 19 individual data sources) of cancer classes for which tobacco smoking increases risk (Table S1). 2,490 were reported to be from tobacco smokers and 1,063 from lifelong non-smokers (Table 1) enabling investigation of the mutational consequences of smoking by comparing somatic mutations in smokers with those in non-smokers for lung cancer (small cell, squamous and adenocarcinoma), larynx, pharynx, oral cavity, esophagus (squamous and adenocarcinoma), bladder, liver, cervix, kidney and pancreas cancers (Fig. 1 and Table S2).

**Fig. 1.**
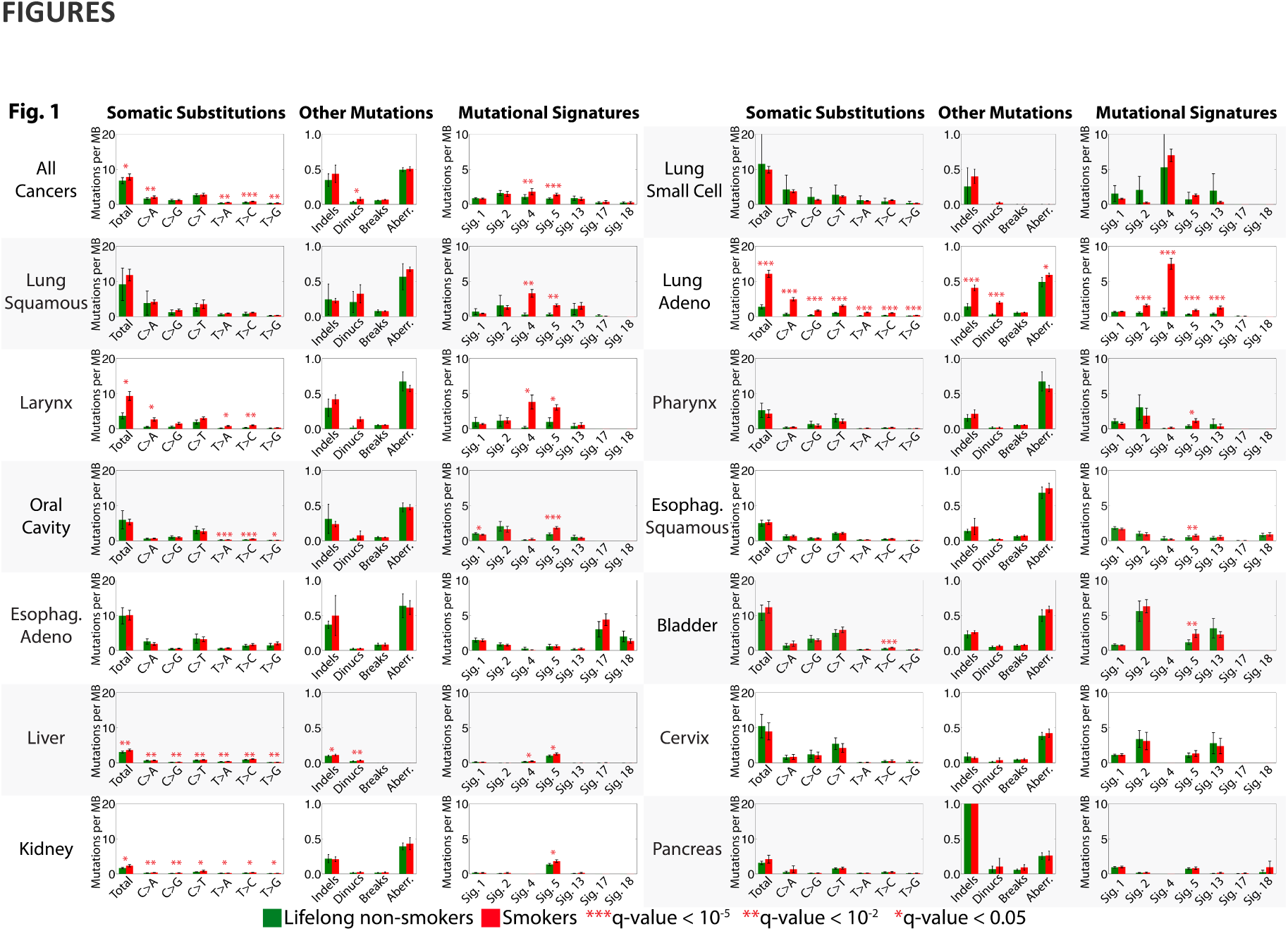
Comparison between tobacco smokers and lifelong non-smokers. Bars are used to display average values for numbers of somatic substitutions per megabase, numbers of indels per megabase, numbers of dinucleotide mutations per megabase, numbers of breakpoints per megabase, fraction of the genome that shows copy number changes and numbers of mutations per megabase attributed to mutational signatures found in multiple cancer types associated with tobacco smoking. Green bars are non-smokers, while red bars are smokers. Comparisons between smokers and non-smokers for all features, including mutational signatures specific for a cancer type and overall DNA methylation are provided in Table S2. Error bars correspond to 95% confidence intervals for each feature. Each q-value is based on a two-sample Kolmogorov-Smirnov test corrected for multiple hypothesis testing for all features in a cancer type. Cancer types are ordered based on their age adjusted odds ratios for smoking as provided in Table 1. Data for numbers of breakpoints per megabase and fraction of the genome that shows copy number changes were not available for liver cancer and small cell lung cancer. Adeno stands for Adenocarcinoma; Esophag. stands for Esophagus.

We first compared total numbers of base substitutions, small insertions and deletions (indels) and genomic rearrangements. Total base substitutions were higher in smokers compared to non-smokers for all cancer types together (1.2-fold, q-value=2.2 × 10^−2^ from two-sample Kolmogorov-Smirnov test corrected for multiple hypothesis testing) and for individual cancer types in lung adenocarcinoma (4.6-fold, q-value=1.2 × 10^−30^), larynx (2.5-fold, q-value=3.7 × 10^−2^), liver (1.2-fold, q-value=1.2 × 10^−4^) and kidney cancers (1.4-fold, q-value=1.4 × 10^−2^). Total numbers of indels were higher in smokers compared to non-smokers in lung adenocarcinoma (2.9-fold, q-value=1.7 × 10^−13^) and liver cancer (1.1-fold, q-value=1.1 × 10^−2^). There were few whole genome sequenced cases and thus it was not possible to directly compare genomic rearrangements between smokers and non-smokers, except in pancreatic and liver cancer where there were no statistically significant differences (Table S2). However, sub-chromosomal copy number changes entail genomic rearrangement and thus serve as surrogates for rearrangements. Smokers with lung adenocarcinoma showed more copy number aberrations (1.2-fold, q-value=1.2 × 10^−2^) than non-smokers (Table S2). No difference was found in any other tobacco associated cancer type (Table S2).

We then extracted mutational signatures, estimated the contributions of each mutational signature to each cancer genome and compared the numbers of mutations attributable to each signature in smokers and non-smokers. Differences between smokers and non-smokers were seen for signatures 2, 4, 5, 13 and 16 (the mutational signature nomenclature is that used in COSMIC, http://cancer.sanger.ac.uk/cosmic/signatures, and in previous reports *(16–18))*.

Signature 4 is characterized predominantly by C>A mutations with smaller contributions from other base substitution classes (Fig. 2 and Fig. S1). It was found only in cancer types in which tobacco smoking increases risk and almost exclusively in those derived from epithelia directly exposed to tobacco smoke, including lung adenocarcinoma, squamous and small cell carcinomas and larynx, pharynx, oral cavity and esophagus cancers, but also in liver cancer (Fig. S2 and S3). Signature 4 is very similar to the mutational signature induced *in vitro* by exposing cells to the carcinogen benzo[*a*]pyrene (cosine similarity=0.94, where 1.00 is a perfect match; Fig. 2 and Fig. S3), a constituent of tobacco smoke *(19)*. The similarity extends to the presence of a transcriptional strand bias indicative of transcription-coupled nucleotide excision repair of bulky DNA adducts on guanine (Fig. S1), the proposed mechanism of DNA damage by benzo[*a*]pyrene and other mutagenic tobacco smoke constituents. Thus, signature 4 is likely the direct mutational consequence of misreplication of DNA damage (adducted bases) induced by tobacco smoke carcinogens.

**Fig. 2.**
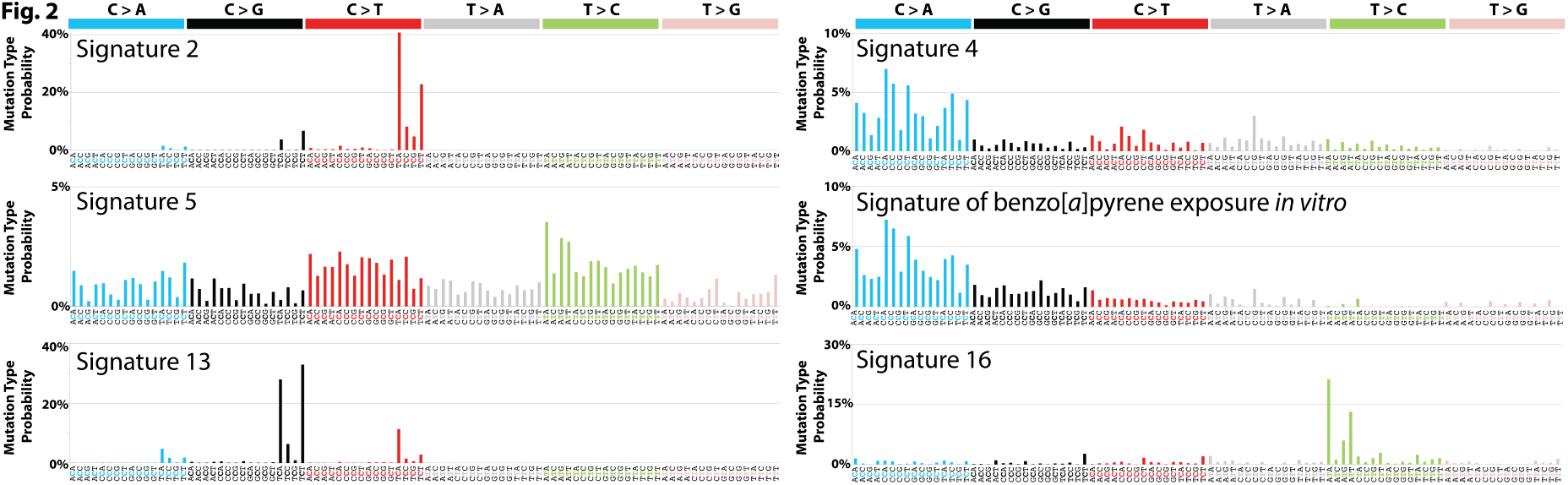
Patterns of mutational signatures associated with tobacco smoking. Signatures are depicted using a 96 substitution classification defined by the substitution type and sequence context immediately 5′ and 3′ to the mutated base. Different colors are used to display different types of substitutions. The percentages of mutations attributed to specific substitution types are on the vertical axes, while the horizontal axes display different types of substitutions. Mutational signatures are depicted based on the trinucleotide frequency of the whole human genome. Signatures 2, 4, 5, 13 and 16 are extracted from cancers associated with tobacco smoking. The signature of benzo[*a*]pyrene is based on *in vitro* experimental data *(19)*. Numerical values for these mutational signatures are provided in Table S6.

Most lung (adenocarcinoma, squamous, small cell) and larynx cancers from smokers showed a large signature 4 mutation burden (Fig. 3). There were greater numbers of signature 4 mutations in cancers from smokers compared to non-smokers in all cancer types together (1.7-fold, q-value=5.8 × 10^−5^) and the difference was substantial in lung squamous (12.4-fold, q-value=3.0 × 10^−3^), lung adenocarcinoma (9.8-fold, q-value=3.9 × 10^−36^) and larynx cancers (20.3-fold, q-value=2.3 × 10^−2^) accounting, in large part, for differences in total numbers of base substitutions (data from only three non-smokers were available for small cell lung cancer; Table 1). 13.8% of lung cancers from non-smokers showed large numbers of signature 4 mutations (>1 mutation per MB) and may be due to passive smoking, misreporting of smoking habits or errors in annotation (Fig. 3). Signature 4 mutations were also present in cancers of the oral cavity, pharynx and esophagus, albeit in much smaller numbers. This may be related to a lesser degree of direct access by tobacco smoke and/or more efficient clearance. Differences in the mutation burden attributed to signature 4 between smokers and non-smokers were not observed for these cancers (Fig. 1 and Table S2). Signature 4 mutations were similarly found at low levels in cancers of the liver, an organ not directly exposed to tobacco smoke, and were elevated in smokers compared to non-smokers (1.4-fold, q-value=1.8 × 10^−2^).

**Fig. 3.**
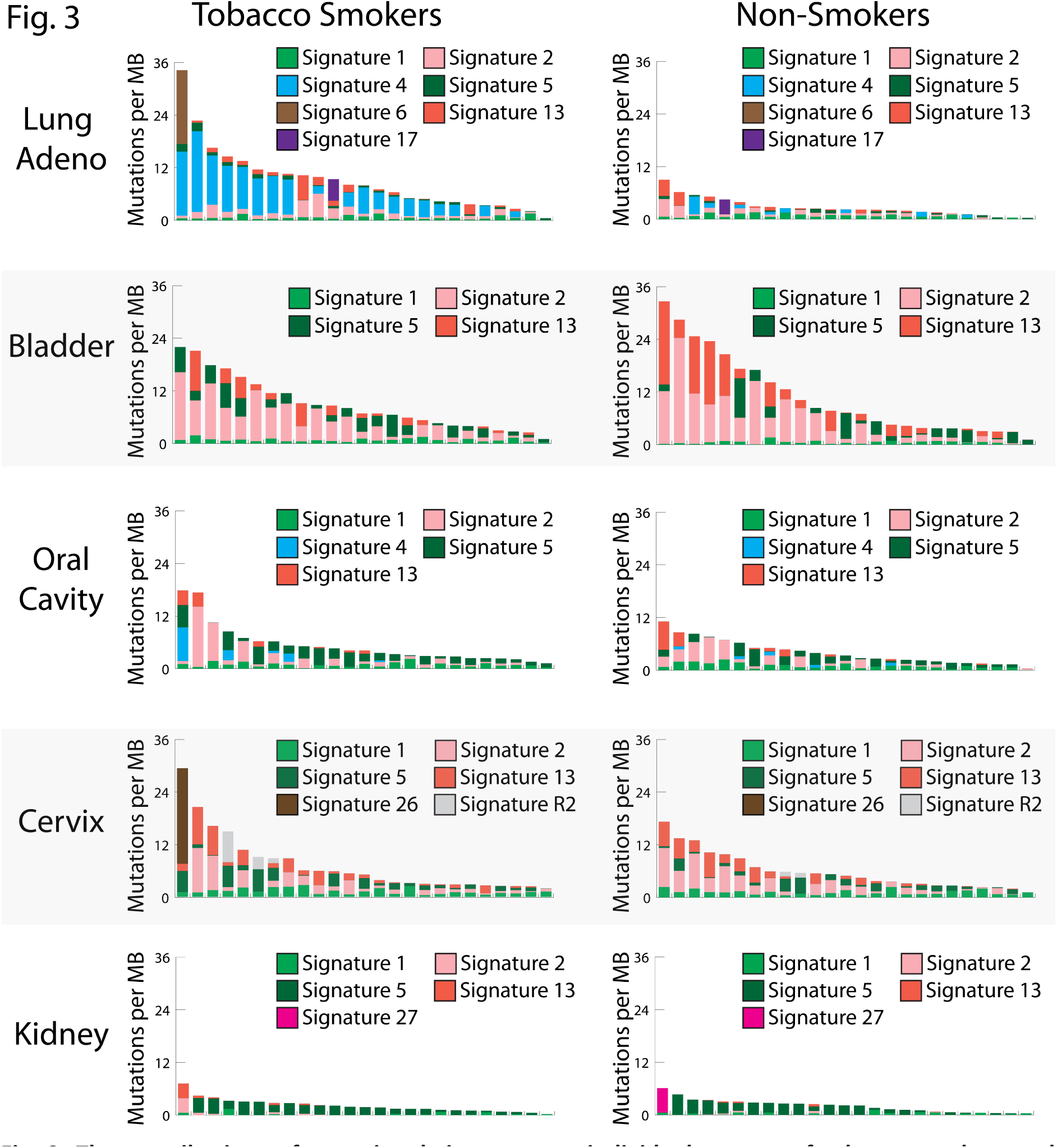
The contributions of mutational signatures to individual cancers of tobacco smokers and non-smokers of selected cancer types. Each panel contains 25 randomly selected cancer genomes (represented by individual bars) from either smokers or non-smokers in a given cancer type. The y-axes reflect numbers of somatic mutations per megabase. Each bar is colored proportionately to the numbers of mutations per megabase attributed to the mutational signatures found in that sample. Naming of mutational signatures is consistent with COSMIC, http://cancer.sanger.ac.uk/cosmic/signatures, and with previous reports *(16–18)*.

Signature 4 was not extracted at all from cancers of the bladder, cervix, kidney or pancreas, despite the known risks for these cancer types conferred by smoking and the presence of many smokers in these series. It was also not extracted from cancers of the stomach, colorectum, ovary and acute myeloid leukemia for which we do not know the smoking status in the analyzed series but among which many cases are likely to have been smokers. The tissues from which all these cancer types are derived are not directly exposed to tobacco smoke. We performed simulations to evaluate whether the absence of signature 4 in these cancer types is due to statistical limitations. Between 1 and 1,000 mutations due to signature 4 were *in silico* added to the mutational catalogues of each sample from bladder, cervix and kidney cancers (Supplementary Text; Fig. S4). As few as 10 somatic mutations, reflecting approximately 0.3 mutations per megabase in an exome, were sufficient to report the presence of signature 4 in more than 25% of the samples in the three cancer types indicating that signature 4 would have been detected if present at these levels. The absence of signature 4 suggests that misreplication of direct DNA damage due to tobacco smoke constituents does not contribute substantially to mutation burden in these cancer types even though studies of DNA adducts have indicated that tobacco-associated DNA damage is present in the tissues from which they arise *(7)*.

Signatures 2 and 13 are characterized predominantly by C>T and C>G mutations respectively at T<u>C</u>N trinucleotides (the mutated base is underlined) and have been attributed to overactive DNA editing by APOBEC cytosine deaminases *(20)*. The cause of the over-activity in most cancers has not been established although the enzyme family is implicated in the cellular response to entrance of foreign DNA, retrotransposon movement and local inflammation *(21)*. Signatures 2 and 13 showed more mutations in smokers than non-smokers in lung adenocarcinoma (3.0-fold with q-value=5.9 × 10 ^11^ for signature 2, and 3.4-fold with q-value=7.8 × 10^−12^ for signature 13). Since they are found in many other cancer types, where they are apparently unrelated to tobacco smoking, it seems unlikely that the signatures 2 and 13 mutations associated with smoking in lung adenocarcinoma are direct consequences of misreplication of DNA damage induced by tobacco smoke constituents. More plausibly, the cellular machinery underlying signatures 2 and 13 is activated by tobacco smoke, perhaps as a result of inflammation arising from deposition of particulate matter.

Signature 5 is characterized by mutations distributed across all 96 subtypes of base substitution, with predominance of T>C and C>T mutations (Fig. 2) and evidence of transcriptional strand bias for T>C mutations *(18)*. Signature 5 is found in all cancer types, including those unrelated to tobacco smoking, and in most cancer samples. It is “clock-like” in that the number of mutations attributable to this signature correlates with age of diagnosis in many cancer types *(17)*. Signature 5, together with signature 1, is thought to contribute to mutation accumulation in most normal somatic cells and in the germline *(17, 22)*. The mechanism underlying signature 5 is not well understood, although signature 5 mutations have recently been shown to be enriched in bladder cancers harboring inactivating mutations in *ERCC2* which encodes a component of the nucleotide excision repair machinery *(23)*.

Signature 5 mutations were increased in smokers compared to non-smokers in all cancer types together (1.7-fold, q-value=2.1 × 10^−6^), and in lung squamous (5.1-fold, q-value=5.2 × 10^−3^), lung adenocarcinoma (3.0-fold, q-value=1.0 × 10^−12^), larynx (3.1-fold, q-value=3.7 × 10^−2^), pharynx (2.8-fold, q-value=2.7 × 10^−2^), oral cavity (2.0-fold, q-value=8.4 × 10^−8^), esophageal squamous (1.5 fold, q-value= 1.3 × 10^−3^), bladder (2.0-fold, q-value=3.3 × 10^−4^), liver (1.3-fold, q=3.3 × 10^−2^) and kidney (1.4-fold, q=1.4 × 10^−2^) cancers (Table S2). The association of smoking with signature 5 mutations across these nine cancer types therefore includes some for which the risks conferred by smoking are comparatively modest and normal progenitor cells which are not directly exposed to cigarette smoke (Table 1). Given the widespread presence of signature 5 in non-smokers, it seems unlikely that signature 5 mutations associated with tobacco smoking are direct consequences of misreplication of DNA damaged by tobacco carcinogens. Rather, it is more plausible that smoking affects the machinery generating signature 5 mutations *(23)*.

Signature 16 is predominantly characterized by T>C mutations at A<u>T</u>N trinucleotides (Fig. 2), exhibits a strong transcriptional strand bias consistent with almost all damage occurring on adenine (Fig. S5) and has only been found thus far in liver cancer. The underlying mutational process is currently unknown. Signature 16 exhibited a higher mutation burden in smokers compared to non-smokers in liver cancer (1.8-fold, q-value=1.0 × 10^−2^).

For smokers with lung (small cell, squamous and adenocarcinoma), larynx, pharynx, oral cavity, esophageal (squamous and adenocarcinoma), bladder, liver, cervix, kidney and pancreas cancers quantitative data on cumulative exposure to tobacco smoke were available (Table S1). Total numbers of base substitution mutations positively correlated with pack years smoked for all cancer types together (rate=50.0 somatic substitutions accumulated per genome per pack year smoked, q-value=3.7 × 10^−5^from robust linear regression analysis corrected for multiple hypothesis testing) and for lung adenocarcinoma (rate=150.5, q-value=1.4 × 10^−3^; Table S3). For individual mutational signatures, correlations with pack years smoked were found in multiple cancer types for signatures 4 and 5. Signature 4 correlated with pack years in lung squamous (rate=51.6, p-value=2.2 × 10^−2^), lung adenocarcinoma (rate=93.7, p-value=1.3 × 10^−2^), larynx (rate=97.2, p-value=2.7 × 10^−2^) and liver cancer (rate=6.4, p-value=3.9 × 10^−2^). Signature 5 correlated with pack years in all cancers together (rate=16.3, q-value=2.6 × 10^−9^), in lung adenocarcinoma (rate=6.4, q-value=3.2 × 10^−2^), in pharynx (rate=38.5, q-value=2.7 × 10^−2^), in oral cavity (rate=22.7, q-value=5.1 × 10^−3^) and in bladder (rate=18.3, q-value=4.6 × 10^−2^) cancers (Table S3). In lung adenocarcinoma, correlation with pack years smoked was also observed for signature 2 (12.9, q-value=3.1 × 10^−2^) and signature 13 (13.9, q-value=6.8x 10^−3^).

Consistent with these results, previous studies have reported that the total number of substitutions is higher in smokers compared to non-smokers in lung adenocarcinoma, mainly due to C>A substitutions *(24, 25)*, and that mutations attributed to signatures 4 and 5 are elevated in smokers with lung adenocarcinoma *(18)*. Recent reports have shown a smoking associated elevation of signature 4 in liver cancer *(26)* and elevation of signature 5 in bladder cancer *(23)*.

Overall levels of CpG methylation of DNA from cancer tissues were similar in smokers and non-smokers for all examined cancer types (Fig. S6). Interrogation of 470,000 CpGs by methylation arrays revealed that individual CpGs were differentially methylated in smokers compared to non-smokers only in lung adenocarcinoma and oral cancer (>5% difference in methylation that satisfies a Bonferroni threshold of 10^−7^). 369 CpGs were hypomethylated and 65 were hypermethylated in lung adenocarcinoma, with five hypomethylated and three hypermethylated in oral cancer (Fig. S7). CpGs exhibiting differences in methylation were neither associated with known cancer genes more than expected by chance nor with genes previously shown to be hypomethylated in normal blood or buccal cells of tobacco smokers (Fig. S8; Tables S4 and S5) *(27)*. The remaining cancer types showed no individual CpGs with significant methylation differences between smokers and non-smokers (Fig. S7). In lung adenocarcinoma and in pharyngeal cancers, the inter-sample variability of methylation levels of CpGs, measured using the inter-quartile range, had more than 20% average increase in smokers compared to non-smokers (Fig. S9).

The genomes of tobacco smoking-associated cancers permit reassessment of our understanding of how tobacco smoke causes cancer. Consistent with the proposition that an increased mutation load caused by tobacco smoke contributes to increased cancer risk, the total somatic mutation burden is elevated in smokers compared to non-smokers in lung adenocarcinoma, larynx, liver and kidney cancers. However, differences in total mutation burden were not observed in the other tobacco smoking-associated cancer types and in some we did not find any statistically significant smoking-associated differences in mutation load, mutational signatures or DNA methylation. Caution should be exercised in the interpretation of the latter observations. In addition to limitations of statistical power, multiple rounds of clonal expansion over many years are often required before a neoplasm becomes a symptomatic cancer. It is thus conceivable that, in the normal tissues from which smoking associated cancer types originate, there are more somatic mutations in smokers than in non-smokers (or differences in methylation) but that these differences become obscured during the intervening clonal evolution. Moreover, some theoretical models predict that relatively small differences in mutation burden caused by tobacco smoking in pre-neoplastic cells could account for the observed increases in cancer risks *(28)* and others that differences in mutation burden between smokers and non-smokers need not necessarily be observed in the final cancers (Supplementary Text and Fig. S6). Thus, increased somatic mutation loads in precancerous tissues may still explain the smoking-induced risks of most cancer types, although other mechanisms have been proposed *(29, 30)*.

The generation of the increased somatic mutation burden by tobacco smoking appears, however, to be relatively complex. Smoking is correlated with increases in base substitutions of multiple different mutational signatures, together with increases in small indels and copy number changes. The extent to which these distinct mutational processes operate differs between tissue types, at least in part depending on the degree of direct exposure to tobacco smoke, and their mechanisms range from misreplication of DNA damage caused by tobacco smoke constituents to activation of more generally operative mutational processes.

## ACKNOWLEDGEMENTS

This work was supported by the Wellcome Trust (grant number 098051). S.N.-Z. is a Wellcome-Beit Prize Fellow and is supported through a Wellcome Trust Intermediate Fellowship (grant WT100183MA). P.J.C. is personally funded through a Wellcome Trust Senior Clinical Research Fellowship (grant WT088340MA). L.B.A. is personally supported through a J. Robert Oppenheimer Fellowship at Los Alamos National Laboratory. This research used resources provided by the Los Alamos National Laboratory Institutional Computing Program, which is supported by the U.S. Department of Energy National Nuclear Security Administration under Contract No. DE-AC52-06NA25396. Research performed at Los Alamos National Laboratory was carried out under the auspices of the National Nuclear Security Administration of the United States Department of Energy. This work was supported by the Francis Crick Institute which receives its core funding from Cancer Research UK, the UK Medical Research Council, and the Wellcome Trust. D.H.P. is funded by Cancer Research UK (grant number C313/A14329). P.V. was partially supported by the project EXPOSOMICS, grant agreement 308610-FP7 (European Commission). Y.T. and T.S. are supported by the Practical Research for Innovative Cancer Control from Japan Agency for Medical Research and Development (15ck0106094h0002), and National Cancer Center Research and Development Funds (26-A-5). We would like to thank The Cancer Genome Atlas (TCGA), the International Cancer Genome Consortium (ICGC), and the authors of all studies cited in Tables S1 for providing free access to their somatic mutational data.

## LIST OF SUPPLEMENTARY MATERIALS

**Methods**

**Supplementary Text**

**Fig. S1.** Comparison of transcriptional strand bias between *in vitro* exposure to benzo[*a*]pyrene and signature 4 extracted from all tobacco-associated cancer types.

**Fig. S2.** Extracting the pattern of signature 4 from different cancer types.

**Fig. S3.** Similarity between extraction of signature 4 across different cancer types and other known mutational signatures.

**Fig. S4.** Sensitivity for detecting signature 4 in cervix, bladder and renal cancer.

**Fig. S5.** Strong transcriptional strand bias of signature 16.

**Fig. S6.** Comparison between lifelong non-smokers and smokers based on overall CpG methylation profiles.

**Fig. S7.** Differentially methylated individual CpGs in tobacco smokers across cancers associated with tobacco smoking.

**Fig. S8.** Cancer tissue methylation of individual CpGs near genes differentially methylated in blood or buccal cells of smokers.

**Fig. S9.** Interquartile range of methylation levels in tobacco-associated cancer types.

**Fig. S10.** Mutation rate variation across normal cells can lead to complex relationships between cancer mutation burden and cancer risk.

**References and notes *(31*–*52)***

## Author Contributions

**Table S1:** Detailed information about each examined tobacco-associated cancer sample.

**Table S2:** Comparison of features of tobacco smokers to the ones of lifelong non-smokers.

**Table S3:** Relationships between mutational signatures and pack years smoked.

**Table S4:** Individual CpGs with differential methylation in lung adenocarcinoma.

**Table S5:** Individual CpGs with differential methylation in oral cancer.

**Table S6:** Numerical patterns of mutational signatures associated with tobacco smoking.

